# Dynamical origins of heat capacity changes in enzyme catalysed reactions

**DOI:** 10.1101/165324

**Authors:** Marc W van der Kamp, Erica J. Prentice, Kirsty L. Kraakmann, Michael Connolly, Adrian J. Mulholland, Vickery L. Arcus

## Abstract

Heat capacity changes are emerging as essential for explaining the temperature dependence of enzyme-catalysed reaction rates. This has important implications for enzyme kinetics, thermoadaptation and evolution, but the physical basis of these heat capacity changes is unknown. Here we show by a combination of experiment and simulation, for two quite distinct enzymes (dimeric ketosteroid isomerase and monomeric alpha-glucosidase), that the activation heat capacity change for the catalysed reaction can be predicted through atomistic molecular dynamics simulations. The simulations reveal subtle and surprising underlying dynamical changes: tightening of loops around the active site is observed as expected, but crucially, changes in energetic fluctuations are evident across the whole enzyme including important contributions from oligomeric neighbours and domains distal to the active site. This has general implications for understanding enzyme catalysis and demonstrating a direct connection between functionally important microscopic dynamics and macroscopically measurable quantities.

---

A critical variable for the rate of a reaction is temperature. For uncatalyzed chemical reactions, the rate of reaction typically increases exponentially with increasing temperature, as described by the Arrhenius and Eyring equations^
1,2
^. In reactions catalysed by enzymes, the effects of temperature are complex and include (often opposing) contributions from active site geometry and reactivity, protein stability, conformational changes and temperature-dependent regulation. Changes in temperature can also potentially affect features of the enzyme catalysed reaction outside the chemical step such as substrate binding, product release and conformational changes. Despite these complexities, enzymes generally show a characteristic temperature profile including an optimum temperature (*T_opt_
*) for activity above which rates decline with increasing temperature. The decline in rate above *T_opt_
* cannot simply be explained by enzyme unfolding at higher temperatures and deviations from Eyring behaviour are also often seen at temperatures below *T_opt_
* ^
3-5
^. We recently developed macro-molecular rate theory (MMRT)^
6,7
^, which explains the temperature dependence of enzymes including an intrinsic *T_opt_
* in the absence of denaturation by introducing the concept of heat capacity changes along the reaction coordinate: the heat capacity (*C_p_
*) for the enzyme–substrate complex is generally larger than *C_p_
* for the enzyme-transition state complex, in enzymes for which the chemical reaction is rate limiting. Hence, the activation heat capacity, 
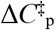
, for the enzyme-catalysed reaction is generally negative (Fig. 1a; in case product release instead of chemical reaction is rate limiting, a small positive heat capacity for reaction is possible^
8
^). We have demonstrated that this accounts for the curvature observed in Eyring plots for a number of enzymes^
6
^.

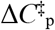
 is a statistical thermodynamic property for the catalysed reaction that describes the difference in heat capacity between the thermodynamic ensemble in the ground state and that at the transition state. It can be determined experimentally^
9
^, and can also be calculated from the variance in enthalpy at equilibrium for each of these states^
10
^:

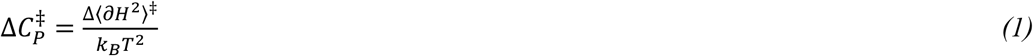

In principle, atomistic molecular dynamics simulations at equilibrium can provide a distribution of enthalpies from which the variance 〈𝜕*H* ^
2
^〉 (the mean squared fluctuation in the enthalpy) may be calculated. To do so, the ensemble for the enzyme-substrate complex and separately, that for the enzyme transition state complex, should be simulated.

Here, we experimentally determine the value for 
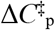
 from the temperature dependence of the rate in the absence of enzyme denaturation for two quite different enzymes: the small, dimeric ketosteroid isomerase (KSI) and the large, monomeric α-glucosidase MalL (Fig. 1). In parallel, we employ extensive MD simulations (10 μs per enzyme) to obtain heat capacity differences between two states along the reaction pathway. KSI is a very well-studied enzyme that is involved in steroid biosynthesis and degradation: it performs two consecutive proton transfers to shift the position of a C=C double bond^
11
^ (Supplementary Fig. 1). MalL is a large α-glucosidase: it hydrolyses terminal non-reducing (1→6)-linked α-glucose residues in a two-step reaction, releasing α-glucose^
12
^ (Supplementary Fig. 2). Previously, we have shown by experiment that there is a large change in heat capacity for this enzyme-catalysed reaction and that single point mutations can dramatically alter the temperature dependence of the rate by altering the heat capacity of either the enzyme-substrate complex or the enzyme-transition state complex^
7
^.

**Figure 1.**
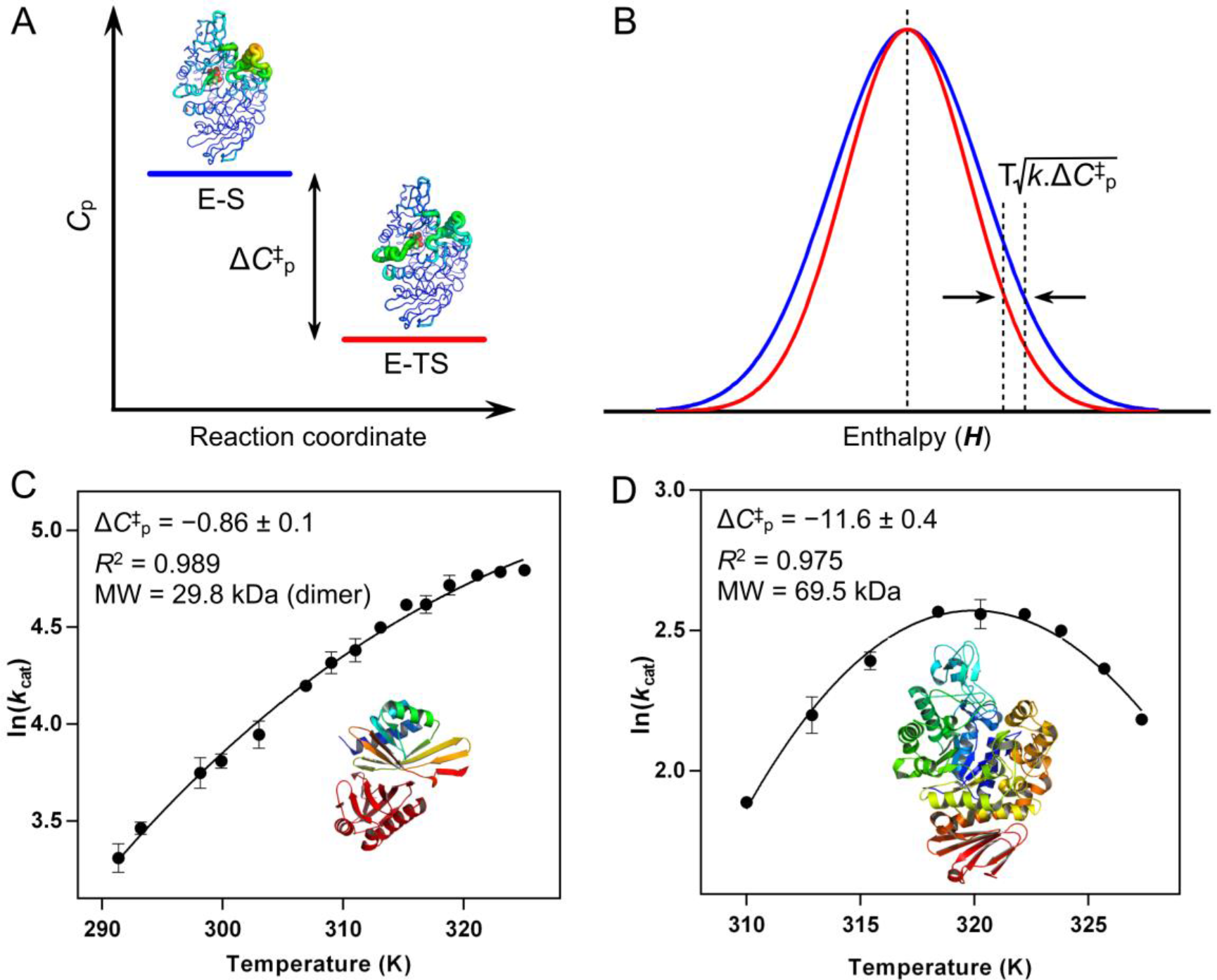
Basis of a negative 
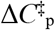
 **and its determination through experiment.** A, Conceptual depiction of a difference in *C*
_p_ between the enzyme-substrate (E-S) and enzymetransition state (E-TS) complexes along a reaction, resulting in a negative 
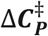
. **B**, Conceptual depiction of differences in enthalpy distribution at the E-TS (red) and the E-S states (blue). Arrows indicate the inflection points (at μ+σ), and the difference defines Δ*C*
_p_ between the two states according to the formula given (see equation (1)). **C-D**, Experimentally determined 
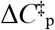
 values (kJ.mol^−1^.K^−1^ ± SE) for the temperature-dependent rates of KSI (**C**) and MalL (**D**). Data is fit with MMRT (see Methods). Error bars, where visible, represent the SD of three replicates. Figures of KSI and MalL are to scale.

**Figure 2.**
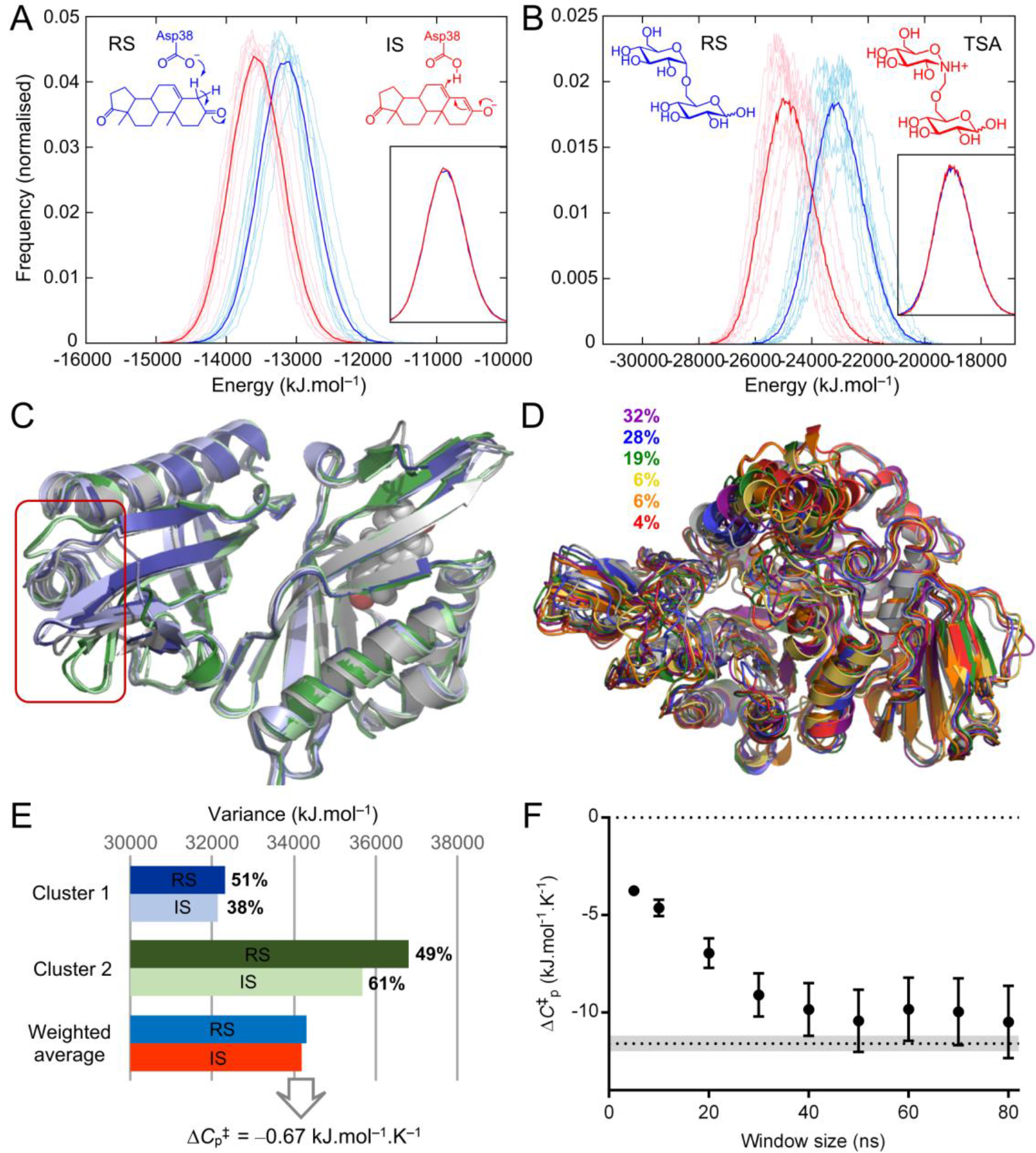
Sampling and 
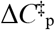
 calculation in simulations. **A-B**, Histograms of energies from 50-500ns MD simulations for KSI (**A**) and MalL (**B**). Thin lines are individual runs, thick lines are the average of 10 runs. Insets show overlay of histograms for both states, and the structures indicate the species simulated (RS – reactant state; IS – intermediate state; TSA – transition state analogue). **C**, Representative structures for the two distinct conformational clusters present in the KSI simulations of both states (reactant state in blue and green, intermediate state in pale blue and green, starting structure in light grey). Box highlights the region with structural differences. **D**, Representative structures for the 6 main conformational clusters in MalL reactant state simulations and their occupancies (starting structure in light grey). **E**, Variance in energies for the two clusters identified in the KSI simulations, with cluster occupancies (in %) and weighted average variance for both states. **F**, Convergence with moving average window size of 
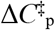
 values calculated for MalL, with value determined from experiment indicated by dotted line (with grey area indicating standard deviation).

Activation heat capacities from simulations and experiment are in good agreement. This shows that prediction of activation heat capacity for enzymes is feasible by simulations, opening a new route to predicting and engineering optimum temperatures for enzyme activities. Further, the simulations provide an atomically detailed picture of the dynamical differences between transition state and Michaelis complexes that gives rise to this behaviour, revealing complex, and intriguing changes in dynamics across the whole enzyme structure. We thus use simulation, for the first time, to interpret the 
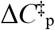
 obtained from macroscopic kinetics measurements in terms of detailed contributions at the atomistic level, providing a link between enzyme structural and energetic molecular fluctuations to its function and thermoadaptation.

## Results

### 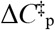 determined by experiment

As shown previously^
6,7
^, 
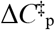
 can be determined by fitting the *ln(rate)*-versus-temperature plot using macromolecular rate theory (MMRT). For MalL, curvature in this plot is very significant (Fig. 1d), and unrelated to unfolding^
5
^. This leads to a negative 
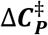
 value of –11.6 ± 0.4 kJ.mol^−1^.K^−1^. For KSI, the curvature is less extreme, but still obvious, leading to a small negative of –0.86 ± 0.1 kJ.mol^−1^.K^−1^ (Fig. 1c). An important consequence of the 
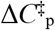
 values of for each enzyme is the position of the optimum temperature (*T_opt_
*) for activity as these parameters are correlated. For example, the large negative 
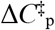
 value for MalL dictates the position of *T_opt_
* at 320 K whereas the much smaller 
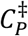
 value for KSI places the *T_opt_
* well above 320 K (in absence of protein unfolding). Similarly, the significant curvature of the MalL temperature dependence means that at lower temperatures, the rate approaches zero much faster for MalL when compared to KSI.

### Heat capacity differences from simulation

Heat capacity differences for enzyme catalyzed reactions can be calculated from Δ〈𝜕*H* ^
2
^〉^‡^ (eq. (1)). To measure Δ〈𝜕*H* ^
2
^〉^‡^ from simulation, there are two main challenges: a) the amount of sampling required for the system to define the enthalpy variance, and b) an accurate and consistent representation of the reactant state (Michaelis complex) and the transition state. A statistical thermodynamic analysis of a 1 ms molecular dynamics (MD) simulation of the bovine pancreatic trypsin inhibitor indicated that 10s of μs of simulation may be needed to converge the heat capacity difference between two conformational states^
13
^. Sampling on the order of (at least) μs is thus expected to be required for reliable identification of heat capacity differences. Such sampling is now routinely feasible with a ‘molecular mechanics’ description of the atoms and their interactions. We thus compare two states, A and B (Fig. 1a), and the difference in heat capacity between these states can be determined by:

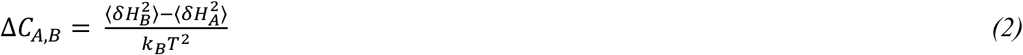

To sample the conformational dynamics of the reactant (E-S or RS) and ‘transition state’ (ETS) enzyme complexes consistently, electronically unstable states (e.g. with half-formed bonds involving enzyme residues) should be avoided for the ‘transition state’ representation. We thus use molecular species that are representative for the transition state (i.e. transition state analogues), and predict that these will show a similar heat capacity change from the reactant state. This prediction has been demonstrated experimentally for human 5’ methylthioadenosine phosphorylase^
9
^. For KSI, a charged enediolate intermediate is formed after the first proton transfer, and this is the key species stabilised by the enzyme for catalysis of the reaction^
14,15
^. We use this intermediate state as a proxy for the two enzyme-transition state complexes (one for each proton transfer) as the intermediate lies between the two transition states at similar energy. The substrate (5–androstene-3,7–dione) and intermediate complexes (Fig. 2a, Supplementary Fig. 1) were built based on KSI in complex with the inhibitor 5α-estran-3,17-dione (PDB 1OHP). For MalL, we obtained an experimental X-ray structure co-crystallised with a stable transition state analogue (Supplementary Table 1; Supplementary Fig. 4) and use this to simulate the thermodynamics of the substrate isomaltose and a close analogue of the transition state species at the rate determining step^
16
^ (Fig. 2b; Supplementary Fig. 2).

A total of 5 μs of MD simulation was run for KSI and MalL in both the substrate-bound and proxy TS representations over ten replicate simulations for each state (Supplementary Results). The force-field potential energy was used as an approximation for the system enthalpy, and was recalculated for the protein-ligand system without explicit water. Considering the *variance* of the enthalpy is the quantity required for 
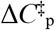
 calculation (eq. (1)) and a *difference* in variance between two states is used to determine 
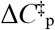
 (eq. (2)) these approximations should be reasonable.

For KSI, the conformational space sampled is limited, with only two distinct structural clusters discernible. The difference between these clusters is in a small region in the unoccupied monomer (Fig. 2c). The *H* variance is significantly different between the clusters, however (Fig. 2e). For 
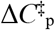
 calculation, we thus calculate the variance of the clusters separately, with the total variance for each state being the average variance weighted by the cluster occupation (Fig. 2e).

MalL samples a larger conformational space than KSI, occupying and regularly switching between a number of structural clusters along the simulation trajectories, related primarily to changes in loops surrounding the active site (Figure 2d; Supplementary Results). Due to the presence of multiple conformational clusters, consideration of the system over the full simulation time overinflates the enthalpy variance. However, calculating variances for each cluster (as for KSI) does not take into account that frequent switches between the distinct conformational states will also contribute to the variance. In addition, several clusters are dominated by one state only (Supplementary Fig. 8). To be independent of clustering and account for switching between conformational substates, enthalpy variance was calculated using a moving window along the simulation trajectory for each simulation, and subsequent averaging. The ‘window’ for the moving average was varied between 5-80 ns and calculated 
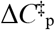
 values converge when the window size is between 40-80 ns (Figure 2f, Supplementary Table 3). Thus, the calculated 
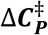
 values for MalL converge on a value of – 10.0 ± 1.7 kJ.mol^−1^.K^−1^ (using a window of 70 ns) which is within the error range of the experimentally determined 
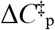
 value of –11.6 ± 0.4 kJ.mol^−1^.K^−1^.

### Local and global contributions to 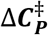

The observation that 
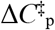
 values calculated from extensive conformational sampling are in agreement with those determined experimentally allows meaningful analysis of the differences between the two ensembles. Striking results emerge from analysing contributions from different parts of the enzymes, by calculating 
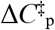
values for parts of the structures (by recalculating energies and their variances for specific regions only; Fig. 3). Energy contributions from interactions with neighbouring regions are not included, and therefore one should not expect these ‘partial’ 
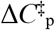
 values to add up to the total value. They do, however, offer new quantitative insights. Conceptually, one may expect differences in partial 
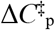
 values to align with regions that differ in flexibility. This is largely true for some small regions with clear differences in flexibility (e.g. residues 46-70 for KSI; residues 250-321 and 374-459 for MalL; see Fig. 3), but is not obvious throughout the structure, especially for larger regions. Crucially, differences in 
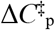
 are distributed across the full protein structure, whereas significant differences in flexibility are limited to regions that interact with the ligand bound in the active site.

**Figure 3.**
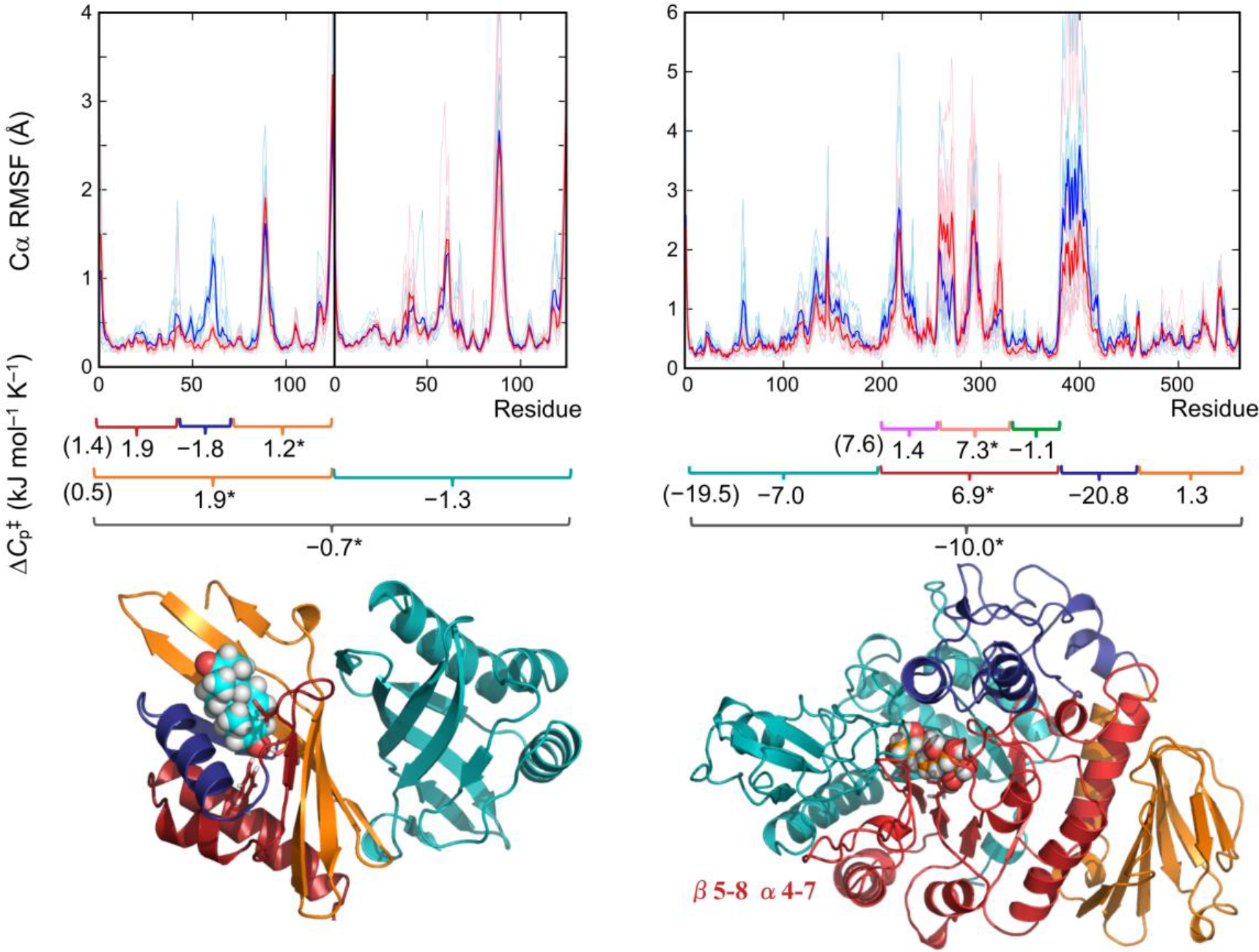
Structural fluctuations and partial variances between reactant state and transition state analogue complexes. Top: root-mean square fluctuations from 50-500ns MD simulations for KSI (left) and MalL (right). Thin lines are individual runs, thick lines the average of 10 runs. Middle: calculated partial 
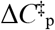
 values for protein regions. Values including contribution from the ligand are indicated (*). Bottom: illustration of KSI (left) and MalL (right) colored based on partial 
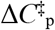
 regions from top pane. Transition-state analogues shown in space-filling spheres.

KSI, as a dimer, offers the opportunity to assess the dynamical role of the monomer that is distal to substrate turnover. The distal monomer of KSI is the main contributor to reduced 
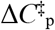
 at the TS (Figure 3A). Overall, the catalytic monomer contributes a positive 
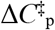
; the N-and C-terminal regions forming the back of the active site and more remote regions contribute a positive 
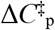
, while helix 48-59 that closes over the active site opposite from the catalytic Asp38 rigidifies and contributes a negative 
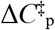
. Implication of the non-catalytic chain as a significant contributor to negative 
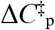
 points to an important role for the oligomer in the temperature dependence of the catalytic process. Enzyme oligomerization is common, indicating an evolutionary advantage^
17
^, however the functional purpose of these quaternary interactions is not well understood. If interactions are optimized to allow global contributions from changes in the distribution of vibrational modes across the multimer^
18
^, oligomerization may provide a means to tune the temperature dependence of rates through global contributions to the overall *C_p_
* change.

The active site of MalL (and TIM barrel enzymes in general^
19
^) sits displaced to one side above the TIM barrel core, interacting with a half of the barrel comprising β5-8 and α4-7 (Figure 3B). Analogous to the ligand bound chain of KSI, the catalytic half of the TIM barrel increases in *C_p_
* at the TS, while more remote protein components including the second TIM half barrel contribute to the overall negative 
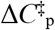
. The lid domain, consisting of a helix-loop-helix extension above the barrel, contributes significantly to the overall reduction in *C_p_
* at the TS, consistent with a role shielding the active site from solvent at the catalytic step. The parallels between the KSI dimer and MalL barrel halves are especially noteworthy in that the TIM barrel is argued to have evolutionary origins as a dimer of (βα)_4_ units^
19,20
^, the dynamical origins of which may still be discernible in the now fused structure.

Overall, these data indicate that the decrease of *C*
_P_ between the enzyme-substrate and enzyme-TS complexes is not just a function of rigidification of elements around the active site, but significant contributions are also made by regions remote from the active site, including oligomeric partners.

## Discussion

In enzyme catalysis, 
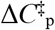
 is emerging as a critical parameter for describing the temperature dependence of enzymatic rates, and as a consequence, for thermal adaptation in enzyme evolution^
6,7
^ The capacity to predictably manipulate enzyme activity with temperature continues to be a sought-after goal in biotechnology^
21
^, but a lack of understanding of the principles governing thermal activity hampers the guided development of enzymes. The *in silico* replication of experimental 
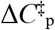
 values gives, for the first time, insight into the atomic-level details of *C*
_P_ changes along the reaction coordinate which govern the temperature dependence of enzyme rates. In turn, this provides a route to engineer temperature optima of enzymes: modifications in enzyme structure that change 
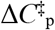
 can be proposed and tested. Due to the difficulty of converging the difference in enthalpy variance that underlies 
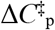
, one cannot expect perfect quantitative agreement between simulation and experiment, but trends and, importantly, atomistic mechanistic details can be extracted.

In two distinct enzyme systems, contributions to reduced 
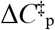
 at the TS come from small domains surrounding the active site as well as domains distant from the catalytic centre. Rigidification of active site loops is expected as these regions may tighten around the transition state ensemble during catalysis. Unexpectedly, domains distal to the active site contribute significantly to the overall negative 
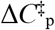
, offsetting positive contributions to 
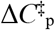
 around the active site (excluding the loops). This observation has implications for the biological importance of both enzyme mass and oligomerization. Previously, enzyme mass has been found to be correlated to catalytic efficiency, leading to the suggestion that vibrational modes (available through the large size of enzymes) act as an energy reservoir, a portion of which is available for catalysis^
6
^. Distal contributions to negative values of 
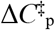
 seen here provide evidence that the necessary changes in the distribution of vibrational modes that give rise to a negative 
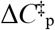
 are dispersed throughout the protein mass. Further, extension of this idea into auxiliary domains (MalL) and dimeric units (KSI) not directly involved with the reaction chemistry suggest that this ‘energy reservoir’ extends further than the catalytic domain. This assigns a significant functional role to distal domains regardless of proximity to the active site, suggesting a functional reason driving the evolution of these domains and interactions.

## Methods

*Cloning, protein expression, purification, and characterization*. Cloning, expression, purification and activity assays of MalL were as described previously^
7
^. Co-crystallization of MalL with 0.5 mM 1-deoxynojirimycin was performed using hanging drop vapor diffusion at 18 °C. Crystals were flash cooled with cryoprotectant comprising of the crystallization mixture with 20 % glycerol for collection at the Australian Synchrotron (MX1). Molecular replacement was performed with the WT MalL apo structure (PDB 4M56)^
7
^ as the search model.

The KSI sequence (*Pseudomonas testosteroni*) with a C-terminal hexa-His tag was optimized for expression in *E. coli*. Expression was carried out over ~24 hours in Luria-Bertani broth at 28 °C. Purified KSI was obtained by a two-step immobilized metal affinity (IMAC)-gel filtration chromatography process. KSI activity was measured *in vitro* using a continuous enzyme assay following the isomerization of 19-nor-androst-5(10)-ene-3,17-dione at 248 nm in phosphate buffer (pH 7.0) for minimal pH change with temperature.

*Experimental 
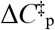
 determination*. Temperature vs. rate profiles were determined by measuring rates in a continuous assay at temperature intervals of 2-4 °C at saturating substrate concentrations. Temperature was controlled via a ThermoSpectronic single cell peltier, and independently checked before and after assays by thermocouple. Initial rates were measured over a period of 10 seconds to limit the effect of denaturation, if present, at elevated temperatures. Temperature profiles were fit with eq. 3 with reference temperature (*T*
_0_) set to *T*
_opt_ – 4:

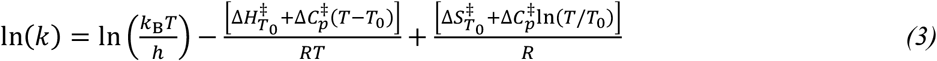

where *k* = rate; *k*
_B_ = Boltzmann constant; *h* = Planks constant; 
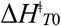
= Enthalpy at *T*
_0_; R = ideal gas constant; 
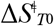
 = entropy at *T*
_0_. The transmission coefficient, κ, is assumed to be 1 for simplicity and is not included.

*MD simulation and analysis*. All simulations and analyses were performed using the Amber package and the ff99SB-ILDN protein force field. For KSI, PDB entry 1OHP was used with Asn38 mutated back to the wild-type Asp and either the substrate or intermediate of the KSI reaction (Fig. 2A) modelled in chain A (based on the co-crystallised inhibitor 5α-estran-3,17-dione); chain B was left empty. General Amber FF (GAFF) parameters with charges from HF/6-31G(d) RESP fitting (RED server: http://upjv.q4md-forcefieldtools.org/REDS/) were used for the ligand. Asp38 was treated as protonated only for the intermediate state in chain A. Asp99 was protonated in both chains, with all other ionizable residues in their standard states. All three Histidines were singly protonated on Nε2 and some Asn/His residue side-chains were rotated by 180° to obtain an optimal hydrogen bond network. For MalL, chain A from the 1-deoxynojirimycin bound structure obtained here (PDB entry 5WCZ) was used with missing atoms built in with COOT based on electron density where available. The substrate isomaltose was placed in the active site by overlay with PDB entry 3AXH (E277A MalL from *S. cerevisiae*; Cα RMSD 0.84 Å) and simulated using GLYCAM (06j-1) parameters. The transition state analogue used in simulation was placed in the active site based on the modelled position of isomaltose and the position of 1-deoxynojirimycin in our co-crystal structure, and simulated using GLYCAM for the glucose unit, and GAFF (with HF/6-31G(d) RESP fitted charges) for the unit containing the protonated nitrogen. Asp63 and Glu371 were treated as protonated in both states, with the catalytic residues Asp199 unprotonated and Glu255 protonated (in line with the mechanism). Other ionizable residues were in their standard protonation states, with His161 singly protonated on Nδ1 and all other His on Nε2. The preparation/equilibration protocol was as follows: solvation in a truncated octahedral of TIP4P(Ew) water molecules (keeping all crystallographic waters) with Na+ ions added to neutralize overall charge (ion positions randomized for each independent run), brief minimization followed by heating in 20ps to 300 K (KSI) or 320 K (MalL) with positional restraints on Cα atoms (5 kcal mol^−1^ Å^−2^), gradual release of restraints in 40 ps, equilibration in the NPT ensemble for 1 ns. 500 ns production simulations were performed in the NVT ensemble with the Berendsen thermostat and loose temperature coupling (10 ps time constant). For both enzymes, restraints were used to maintain the Michaelis complex (with equivalent restraints on the IS or TSA states; Supplementary Methods).

Analysis was performed using 10 ps snapshots from 50-500 ns of the simulations, with force-field energies re-calculated after stripping of solvent and ions. Clustering on the Cα RMSD (excluding the highly flexible C-terminal residues 117-125 in KSI and the N-terminal residues 1-6 in MalL) was performed using the hierarchical agglomerative algorithm using a minimum cluster distance of 1.6 for KSI and 2.1 for MalL. Cα RMSF was calculated using RMSD fitting to a running average coordinates from a time window of 10 ns.

*Data availability*. Coordinates and structure factors for MalL co-crystallised with 1-deoxynojirimycin are deposited in the PDB under accession number 5WCZ. Simulation input files are available from the Dryad Digital Repository (DOI code [to be inserted]).

## Acknowledgments

MWvdK is a BBSRC David Phillips Fellow (BB/M026280/1) and he and AJM thank the BrisSynBio Synthetic Biology Research Centre for funding (BB/L01386X/1). EJP was supported by a University of Waikato Doctoral Scholarship and VLA was supported by the Marsden Fund of New Zealand (08-UOW-057). MC and AJM thank the EPSRC Centre for Doctoral Training in Theory and Modelling in Chemical Sciences (EP/L015722/1).

## Author contributions

MWvdK devised simulation and analysis; MWvdK and EJP performed simulations and analysis, assisted by MC; EJP and KLK performed experiments; MWvdK, EJP, AJM and VLA analysed and interpreted results, and wrote the manuscript. MWvdK and EJP contributed equally to this work.

## References

1 Arrhenius, S. Über die Reaktionsgeschwindigkeit bei der Inversion von Rohrzucker durch Säuren. Z Phys Chem 4, 226-248 (1889).

2 Eyring, H. The activated complex in chemical reactions. J Chem Phys 3, 107-115 (1935).

3 Thomas, T. M. & Scopes, R. K. The effects of temperature on the kinetics and stability of mesophilic and thermophilic 3-phosphoglycerate kinases. Biochemical Journal 330, 1087-1095 (1998).

4 Daniel, R. M. & Danson, M. J. A new understanding of how temperature affects the catalytic activity of enzymes. Trends in Biochemical Sciences 35, 584-591 (2010).

5 Buchanan, C. L. , Connaris, H. , Danson, M. J. , Reeve, C. D. & Hough, D. W. An extremely thermostable aldolase from *Sulfolobus solfataricus* with specificity for non-phosphorylated substrates. Biochemical Journal 3, 563-570 (1999).

6 Arcus, V. L. et al. On the Temperature Dependence of Enzyme-Catalyzed Rates. Biochemistry 55, 1681-1688 (2016).

7 Hobbs, J. K. et al. Change in Heat Capacity for Enzyme Catalysis Determines Temperature Dependence of Enzyme Catalyzed Rates. ACS Chemical Biology 8, 2388–2393 (2013).

8 Nguyen, V. et al. Evolutionary drivers of thermoadaptation in enzyme catalysis. Science 355, 289-293 (2017).

9 Firestone, R. S. , Cameron, S. A. , Karp, J. M. , Arcus, V. L. & Schramm, V. L. Heat capacity changes for transition-state analogue binding and catalysis with human 5′-methylthioadenosine phosphorylase. ACS Chemical Biology 12, 464-473 (2017).

10 Prabhu, N. V. & Sharp, K. A. Heat Capacity in Proteins. Annual Review of Physical Chemistry 56, 521-548 (2005).

11 Ha, N. C. , Choi, G. , Choi, K. Y. & Oh, B. H. Structure and enzymology of Delta5-3-ketosteroid isomerase. Curr Opin Struct Biol 11, 674-678 (2001).

12 Zechel, D. L. & Withers, S. G. Glycosidase Mechanisms: Anatomy of a Finely Tuned Catalyst. Accounts of Chemical Research 33, 11-18 (2000).

13 Fenley, A. T. , Muddana, H. S. & Gilson, M. K. Entropy-enthalpy transduction caused by conformational shifts can obscure the forces driving protein-ligand binding. Proc Natl Acad Sci U S A 109 (2012).

14 van der Kamp, M. W. , Chaudret, R. & Mulholland, A. J. QM/MM modelling of ketosteroid isomerase reactivity indicates that active site closure is integral to catalysis. FEBS Journal 280, 3120-3131 (2013).

15 Fried, S. D. , Bagchi, S. & Boxer, S. G. Extreme electric fields power catalysis in the active site of ketosteroid isomerase. Science 346, 1510-1514 (2014).

16 Cogoli, A. & Semenza, G. A probable oxocarbonium ion in the reaction mechanism of small intestinal sucrase and isomaltase. J Biol Chem 250, 7802-7809 (1975).

17 Ricard, J. & Noat, G. Catalytic efficiency, kinetic co-operativity of oligomeric enzymes and evolution. Journal of Theoretical Biology 123, 431-451 (1986).

18 Kim, T. H. et al. The role of dimer asymmetry and protomer dynamics in enzyme catalysis. Science 355 (2017).

19 Wierenga, R. K. The TIM-barrel fold: a versatile framework for efficient enzymes. FEBS Letters 492, 193-198 (2001).

20 Höcker, B. , Jürgens, C. , Wilmanns, M. & Sterner, R. Stability, catalytic versatility and evolution of the (βα)_8_-barrel fold. Current Opinion in Biotechnology 12, 376-381 (2001).

21 Burton, S. G. , Cowan, D. A. & Woodley, J. M. The search for the ideal biocatalyst. Nat Biotechnol 20, 37-45 (2002).

